# Functional divergence of the repressors of photomorphogenesis SPA2 and SPA3 during Arabidopsis seedling deetiolation

**DOI:** 10.64898/2026.05.21.726765

**Authors:** Zhen Cao, Vivian Feldmann, Ira Trivedi, Ute Hoecker

## Abstract

The COP1/SPA ubiquitin ligase is a key repressor of photomorphogenesis that is inactivated by photoreceptors to initiate light signalling. The four SPA proteins (SPA1-SPA4) confer functional specificity to COP1 during plant growth, yet the underlying molecular mechanisms remain unclear. Here, we used a domain-swap approach in transgenic seedlings to address the functional divergence of SPA2 and SPA3. We show that the respective N-terminal kinase domain determines the contrasting protein stabilities of SPA2 and SPA3 in light-grown seedlings. The instability of SPA2 correlates with a specific ability of the SPA2 N-terminal domain to bind phytochrome A in the light, suggesting that phytochrome A promotes the CUL4^DET1/COP1^-dependent degradation of SPA2 but not of SPA3. We uncover that the coiled-coiled and WD-repeat domains of SPA2 and SPA3 substantially differ in their activity in repression of photomorphogenesis, with those of SPA2 being more active repressors than those of SPA3. Thus, SPA2 combines a potent repressor activity with light-induced instability. We conclude that the evolution of SPA2 instability in response to light counterbalances its inherent strong repressor activity, thereby allowing seedling etiolation in darkness followed by rapid reduction in COP1 activity through SPA2 degradation upon light-exposure as seedlings emerge from soil to initiate photosynthetic growth.

**Highlight:** The repressor of light signaling SPA2 combines a phytochrome A-interacting instability domain with a potent repressor domain to allow greatly contrasting activities of COP1 in skoto- and photomorphogenesis.

## Introduction

As sessile organisms, plants have evolved mechanisms to adapt their growth, development and metabolism to the changing ambient light environment. This includes seed germination, seedling growth, leaf expansion, shade avoidance and flowering time. Seedlings exhibit a particularly strong response to light by opening and greening of their cotyledons and shortening of their hypocotyl (photomorphogenesis, deetiolation). In darkness, in contrast, cotyledons remain closed and pale, while the hypocotyl elongates (skotomorphogenesis, etiolation). To perceive the light, plants harbor several classes of photoreceptors. This includes the type-II phytochromes (phy), such as phyB, which are activated by red light and sense the red/far-red ratio, the type-I phytochrome phyA which senses continuous far-red and blue light, the blue light-perceiving cryptochromes and the UV-B-responsive UVR8 protein (Kami *et al*., 2010). After photo-activation, these photoreceptors initiate light signalling by inactivating a key repressor of photomorphogenesis, the cullin4-based CONSTITUTIVELY PHOTOMORPHOGENIC1/SUPPRESSOR OF PHYA-105 (COP1/SPA) ubiquitin ligase complex. The COP1/SPA complex is thus primarily active in darkness to polyubiquitinate positive regulators of light signalling, mostly transcription factors, which are subsequently degraded in the 26S proteasome. After exposure to light, the inhibition of COP1/SPA activity allows stabilization of these transcription factors, such as HY5 and PAP1, which vastly alter the gene expression program to cause photomorphogenesis (Ponnu and Hoecker, 2021; Zhou and Deng, 2025).

The COP1/SPA complex is likely a tetramer consisting of two COP1 and two SPA proteins of the four-member SPA protein family (SPA1-SPA4). *cop1* and *spa* quadruple mutants undergo constitutive photomorphogenesis, exhibiting a short hypocotyl and open cotyledons in complete darkness. Moreover, they are very dwarfed and flower early in short day photoperiod (Deng *et al*., 1991; Laubinger *et al*., 2004; Ordonez-Herrera *et al*., 2015). This indicates that both COP1 and SPA proteins are important for the activity of COP1/SPA as a repressor of photomorphogenesis. *COP1* is a single-copy gene in Arabidopsis, while the four *SPA* genes have overlapping but also partially distinct functions in light-controlled growth and development (Fittinghoff *et al*., 2006; Laubinger *et al*., 2004; Laubinger *et al*., 2006; Menon *et al*., 2016; Rolauffs *et al*., 2012). The COP1 protein harbors three domains, an N-terminal RING-finger domain, a central coiled-coil domain and a C-terminal WD-repeat domain (Deng *et al*., 1992). All four SPA proteins also carry a C-terminal WD-repeat domain, a central coiled-coil domain, but they differ from COP1 in their N-terminus by carrying an atypical protein kinase domain (Hoecker *et al*., 1999) which was recently shown to exhibit kinase activity (Paik *et al*., 2019). COP1 and SPA proteins homo- and heterodimerize through their coiled-coil domains (Hoecker and Quail, 2001; Saijo *et al*., 2003; Zhu *et al*., 2008). Their WD-repeat domains act as substrate-recognition domains by recognizing a conserved valine-proline (VP) motif present in ubiquitination substrates, though substrates lacking a VP motif are known as well (Lau *et al*., 2019; Ponnu *et al*., 2019; Tao *et al*., 2025; Trimborn *et al*., 2025; Uljon *et al*., 2016). While the WD-repeats of both COP1 and SPA1 bind substrates, they are not fully functionally interchangeable, suggesting that the WD repeat of SPA1 has a function that cannot be taken over by that of COP1 (Kerner *et al*., 2021).

The tight regulation of COP1/SPA activity by light is key to photomorphogenesis. This is achieved by a light-induced interaction of the photoreceptors phytochromes, cryptochromes and UVR8 with the COP1/SPA complex (Favory *et al*., 2009; Holtkotte *et al*., 2017; Lian *et al*., 2011; Sheerin *et al*., 2015; Zuo *et al*., 2011). At least four mechanisms evolved to inhibit COP1/SPA function in response to light. This includes nuclear exclusion of COP1 which is dependent on SPA proteins (Balcerowicz *et al*., 2017; Von Arnim and Deng, 1994), disruption of the COP1-SPA interaction (Lian *et al*., 2011; Lu *et al*., 2015; Sheerin *et al*., 2015), cryptochrome- and UVR8-mediated competitive displacement of substrates from binding to COP1 (Lau *et al*., 2019; Ponnu *et al*., 2019; Trimborn *et al*., 2025) and degradation of SPA1 and SPA2 proteins (Balcerowicz *et al*., 2011).

Multiple COP1/SPA complexes are formed containing different members of the four SPA proteins (Zhu *et al*., 2008), thereby providing specificity to COP1 in the regulation of plant growth and development. A key regulatory mechanism of selective COP1/SPA complexes is the differential stability of the four SPA proteins in light-grown seedlings which greatly impacts light responses. SPA3 and SPA4 proteins are stable in the light, while SPA1 and SPA2 are destabilized in light-exposed seedlings (Balcerowicz *et al*., 2011; Schenk *et al*., 2021). Particularly, SPA2 is very rapidly degraded in response to even very short pulses of light of a very low fluence rate. Light-induced SPA2 degradation is essential for photomorphogenesis because *SPA2* mutations that stabilize a functional SPA2 protein prevent photomorphogenesis even at very high fluence rates. Moreover, the rapid and efficient degradation of SPA2 in the light correlates with the most effective inactivation of SPA2 function in light-grown seedlings when compared to the other three SPA proteins (Chen *et al*., 2015). SPA2 degradation in the light requires the N-terminal kinase domain of SPA2 and COP1, DET1 and CUL4, suggesting that SPA2 is polyubiquitinated by a CUL4^DET1-COP1^-containing E3 ubiquitin ligase (Schenk *et al*., 2021). SPA2 is rapidly degraded in red, far-red as well as in blue light and – most importantly – in a fashion that depends on phytochromes A and B and, even in blue light, is independent of cryptochromes (Chen *et al*., 2016). Thus, light-induced SPA2 degradation provides a phytochrome-specific mechanism of COP1/SPA inactivation.

Taken together, these reports indicate that the evolved distinct protein stabilities of the four SPA proteins are critical for photomorphogenesis. However, the molecular mechanisms responsible for the functional divergence among SPAs are not well understood. We therefore addressed this question by comparing two SPA proteins, the light-stable SPA3 and the light-labile SPA2. Using a domain-swap approach, we show that the N-terminal kinase domain of these SPAs directs the protein stability in light-grown seedlings by affecting the interaction with phytochrome A. Beyond that, our experiments uncover that the C-terminal domains of SPA2 are more effective in repression of photomorphogenesis than those of SPA3. This has important implications on early seedling growth. Thus, *SPA* gene duplications were a driving force to allow divergence of SPA activity for optimal tuning of responses to the ambient light environment.

## Materials and methods

### Plant material, light sources and growth conditions

All genotypes are in the Col-0 background. The genotypes *cop1-4* (Deng and Quail, 1992), *spa1-7 spa2-1 spa3-1* (referred to as *spa123*) (Fittinghoff *et al*., 2006) and *SPA2::SPA2-HA spa123* #32 (Balcerowicz *et al*., 2011) were described previously. All transgenic lines were generated in the *spa1-7 spa2-1 spa3-1* (referred to as *spa123*) background by floral dipping.

LED light sources and growth conditions were described previously. Briefly, seeds were surface-sterilized and plated on MS media without sucrose, incubated at 4°C in darkness for 3 days. To induce germination, plates were subsequently exposed to white light at 21°C for 3 h, transferred back to darkness for approx. 21 h and then kept in darkness, red, far-red or blue light for 3 days at 21°C.

### Hypocotyl length determination

Seedlings were laid flat on the surface of solid MS plates and photographed with a Nikon D5000 digital camera. Hypocotyl length was measured for at least 15 seedlings per genotype and condition using ImageJ1.53 (Wayne Rasband, National Institutes of Health).

### Generation of transgenic Arabidopsis lines

All transgenic lines were generated in the *spa1-7 spa2-1 spa3-1* (referred to as *spa123*) background by floral dipping. Transgenic plants were selected on MS medium containing 15 µg/ml hygromycin. In the T2 generation, lines carrying one transgene insertion site were selected based on a 3:1 segregation for hygromycin resistance : sensitivity. T3 and T4 lines homozygous for the transgene were used for analyses. At least three independent lines were used for each construct.

### Plasmid constructions

The generation of all plasmid constructs is described in the Supplemental materials and methods section.

### Protein isolation from Arabidopsis seedlings

Approximately 0.2 g of seedlings was snap-frozen in liquid nitrogen and ground by an electronic tissue grinder (Retsch MM400; Retsch GmbH, Haan, Germany). The ground powder was resuspended in 300 µl YODA buffer (50 mM Tris/HCl pH 7.5, 150 mM NaCl, 1 mM EDTA, 10% (v/v) glycerol, 0.1% Triton X-100, 1% (v/v) protease inhibitor cocktail (Sigma Aldrich Chemie GmbH, Munich, Germany), 5 mM DTT, 10 µM MG132) and allowed to thaw completely on ice. The mixture was then centrifuged at 13.000 rpm at 4℃ for 20 min and the supernatant was transferred to a new tube. Ten µl of the supernatant was used for a Bradford assay to measure the protein concentration. The remaining supernatant was supplemented with 5 x Laemmli buffer (312.5 mM Tris-HCl pH 6.8, 10% (w/v) SDS, 50% (v/v) Glycerol, 0.25% (w/v) bromophenol blue, 500 mM DTT) and heated at 96℃ for 5 min. The samples were stored at - 20℃.

### Immunodetection of proteins

Protein samples (20 µg) were separated by SDS-PAGE using the Mini PROTEAN Tetra electrophoresis system (Bio-Rad Laboratories, Munich, Germany). Proteins were transferred onto a polyvinylidene difluoride (PVDF) membrane by wet or semi-dry blotting. The membrane was blocked in 1x Roti®-Block solution (Carl Roth, Karlsruhe, Germany) for 1 h at room temperature or overnight at 4 ℃. Membranes were incubated with the primary antibody dissolved in 1 x TBS (20 mM Tris/HCl pH 7.5, 150 mM NaCl) containing 2% non-fat milk powder (VWR International GmbH, Darmstadt, Germany) on a inverting rotator for 1 h at room temperature or overnight at 4 ℃. After incubation, the membranes were washed three times in TBS-T (TBS + 0.1% (v/v) Tween-20) for 10 min each. Subsequently, the membranes were incubated with HRP-coupled secondary antibody at room temperature for 1 h. Afterwards, the membranes were washed twice with 1 x TBS-T for 10 min each and once in 1 x TBS for 10 min. The HRP activity was detected by SuperSignal® West Femto Maximum Sensitivity kit (Thermo Fisher Scientific; Schwerte, Germany). The signal was visualized using the ImageQuantTM LAS 4000 mini-imaging system (GE Healthcare, Piscataway, USA).

### Antibodies

α-HA-HRP (Cat: 12013819001) 1:2000 dilution, Sigma-Aldrich (Munich, Germany); α-HSC70 (Cat: SPA-817) 1:10000, Enzo Life Sciences (Lörrach, Germany); α-Tubulin (Cat: T5168) 1:10000, Sigma-Aldrich (Munich, Germany); α-phyA (Cat: AS07 220) 1:1500, Agrisera (Vännäs, Sweden); α-rabbit IgG-HRP (Cat: AS09 605) Agrisera (Vännäs, Sweden); α-mouse IgG-HRP (Cat: 12-439) 1:50000 Sigma-Aldrich (Munich, Germany).

### Yeast two-hybrid assays

Co-transformation of AD and BD plasmids competent cells was performed using the Frozen-EZ Yeast Transformation II Kit (Zymo Research Europe GmbH, Freiburg, Germany) in accordance with the manufacturer’s instructions. The transformed yeast cells were plated onto SD-L-W dropout plates lacking leucine and tryptophan and incubated at 30 ℃ for 3 days. Ten-15 colonies from each sample were picked from the SD-L-W plates and resuspended in 1 ml sterile ddH2O. The suspensions were vortexed briefly, and two 15 µl spots of each yeast cell suspension were dropped onto fresh SD-L-W plates supplemented with 20 µM phycocyanobilin (Sirius Fine Chemicals SiChem GmbH, Bremen, Germany).. Each combination was prepared in three biological replicates. The plates were incubated in darkness for 24 h at 26℃ followed by exposure to 1 µmol m^−2^s^−1^ red light for 2 days or continued incubation in darkness for 2 additional days. The yeast cells were harvested by scraping from the plates using sterile forceps under either red or green safe light, respectively, resuspended in 1 ml Z-buffer (10.7 g/L Na_2_HPO_4_ x 2 H_2_O, 5.5 g/L NaH_2_PO_4_ x H_2_O, 0.75 g/L KCl, 0.246 g/L MgSO_4_ x 7 H_2_O) and the OD600 was measured. The suspensions were diluted to an OD600 of 1 in a final volume of 300 µl Z-buffer. The samples were subjected to five freeze-thaw cycles in liquid nitrogen to ensure complete cell lysis. Each sample was split into two 150 µl technical replicates, followed by the addition of 700 µl Z-buffer supplemented with 270 µl β-mercaptoethanol/100 ml) . Subsequently, 160 µl Z-buffer containing 4 mg ortho-nitrophenyl-ß-galactoside/ml was added. The samples were incubated at 30 ℃ until a color change to yellow was observed. To terminate the reaction, 400 µl 1M Na_2_CO_3_ was added and the reaction time was recorded. After centrifugation at 13.000 rpm for 10 min, 200 µl of the supernatant was transferred into a 96 well Greiner half-area microplate (Greiner Bio-One International GmbH, Kremsmuenster, Austria) and the absorbance at 420 nm was measured using an infinite RM200 plate reader (Tecan, Männedorf, Switzerland). The β-galactosidase activity was calculated in Miller units according to the yeast protocol handbook (Clontech).

### Luciferase complementation imaging (LCI) assays

The required plasmids expressing the nLUC and cLUC fusion proteins were transformed into the *Agrobacterium tumefaciens* strain GV3101 (pMP90). The Agrobacteria were incubated in 5 ml LB medium supplemented with the appropriate antibiotics and incubated overnight at 28℃ under constant shaking. The overnight culture were harvested by centrifugation at 4000 rpm for 20 min at room temperature. The resulting pellet was resuspended in 2 ml 1 x MES buffer (10 mM MES, 10 mM MgCl_2_,0.2 mM acetosyringone) and the OD600 was measured and adjusted to 0.8 using 1 x MES buffer. The suspensions along with RK19, an anti-silencing strain, were mixed in equal proportions and incubated at room temperature for 20 min to 2 h. The mixture was co-infiltrated into the abaxial side of *N. benthamiana*. After infiltration, N*. benthamiana* plants were maintained in darkness overnight followed by transfer to a greenhouse for 2-3 days under standard growth conditions. For the detection of the luciferase activity, the injected leaves were detached and sprayed with D-luciferin solution (5 mM D-luciferin, 0.025 % Triton X-100). To minimize the chlorophyll autofluorescence, the leaves were incubated in darkness for 5 min before imaging. The luciferase activity was visualized using the ImageQuantTM LAS 4000 mini-imaging system (GE Healthcare, Piscataway, USA).

### Statistical analyses

T-tests were performed using the P values and the number of biological replicates indicated in the figure legends. Each experiment was repeated at least twice.

## Results

### The N-terminal domains of SPA2 and SPA3 carry the information for SPA protein stability in the light to allow optimal seedling deetiolation

The four SPA proteins differ in their protein stability in light-grown seedlings which impacts specificity and key functions. SPA1 and SPA2 are degraded in response to light, while SPA3 and SPA4 remain equally stable in light- and dark-grown seedlings (Balcerowicz *et al*., 2011; Schenk *et al*., 2021). To investigate which domain(s) of the SPA proteins confers the divergent protein stability, we conducted a domain-swap analysis between the highly light-unstable SPA2 protein and the light-stable SPA3 protein. Since deletion of the N-terminal domain of SPA2 strongly stabilizes SPA2 (Schenk *et al*., 2021) we hypothesized that the N-terminal domains of SPA2 and SPA3 may confer distinct information on protein stability. We therefore constructed two domain swaps, fusing the N-terminal domain of SPA2 to the C-terminal coiled-coil and WD-repeat domains of SPA3 (from now on named “DS_233” for domain swap (DS) and the SPA-identity of each of the three domains in the chimeric proteins) and vice versa (DS_322) (Figure 1A). These domain-swap proteins were fused to a 3xHA tag and expressed in transgenic *spa123* triple mutants to be able to compare the protein stability of the chimeric proteins as well as the degree of complementation of the *spa123* mutant phenotype. As controls served lines that expressed non-chimeric SPA2-3xHA (described previously in (Balcerowicz et al., 2011)) and non-chimeric SPA3-3xHA. All transgenes were expressed under the control of the *SPA2* 5’ and 3’ regulatory sequences which allow constitutive, i.e. light-independent expression of the transgenes (Balcerowicz *et al*., 2011; Fittinghoff *et al*., 2006). Therefore, light-dependent changes in SPA-HA protein levels are not corroborated by any effect of light on *SPA-HA* transcript levels.

**Figure 1.**
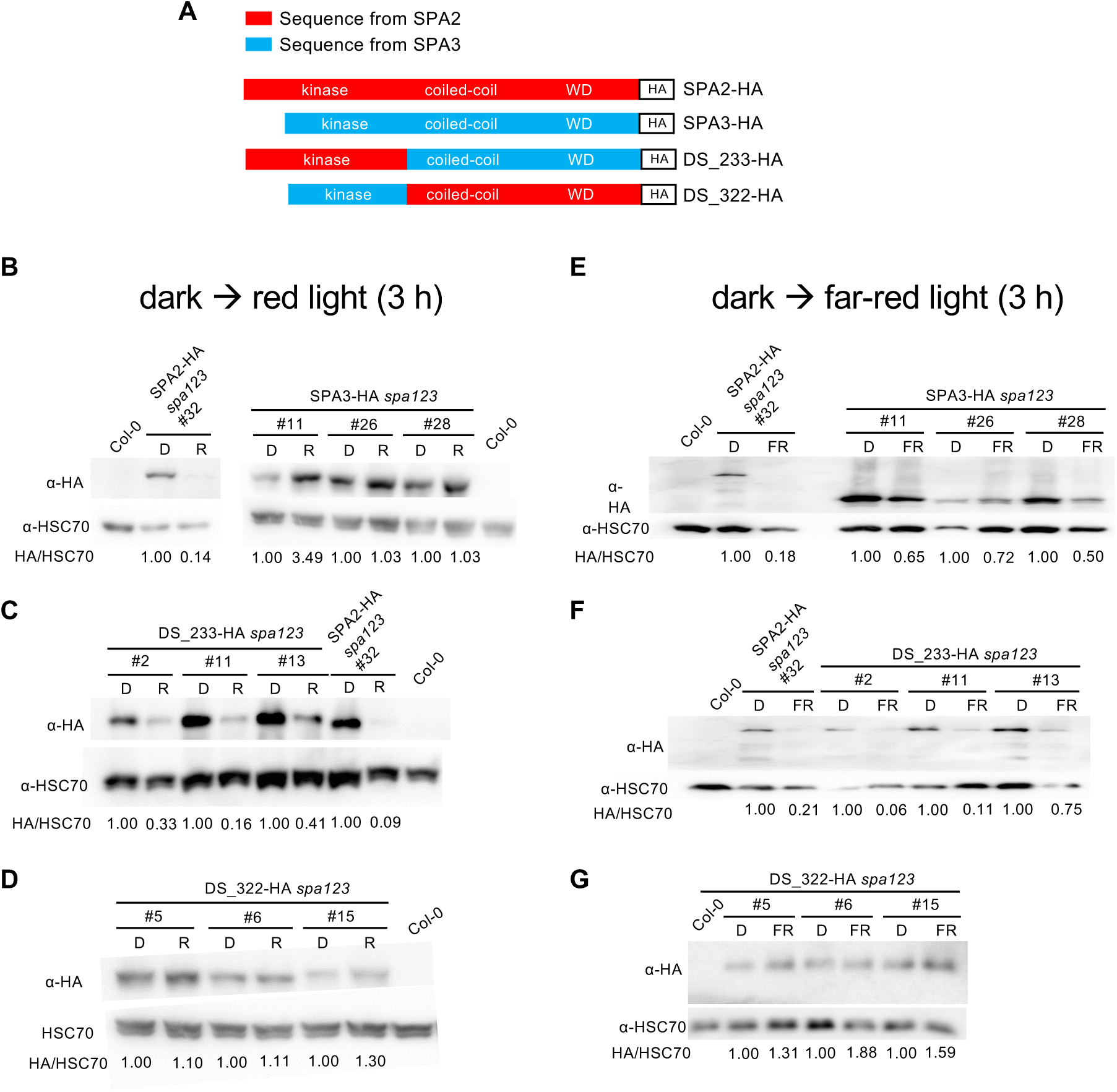
The N-terminal domains of SPA2 and SPA3 confer SPA protein stability information in red and far-red light. A. Schematic representation of domain-swap approach and controls. Full-length SPA2 (SPA2-HA), SPA3 (SPA3-HA) and chimeric domain swap proteins swapping the N-terminal domains between SPA2 and SPA3 (DS_233-HA and DS_322-HA) were expressed under the control of the constitutive *SPA2* 5’ and 3’ regulatory sequences in the *spa123* triple mutant background. The numbers in the domain-swap line names (e.g. 233) refer to the SPA-identity of the respective three domains of SPA proteins: kinase, coiled-coil and WD. All proteins were fused to a triple HA tag (HA). Line numbers reflect independent transgenic lines. B-G. Immunodetection of the indicated HA-tagged SPA proteins in seedlings grown in darkness (D) for 4 days followed by transfer to 1 µmol m^−2^ s^−1^ red light (R) for 3 h (B-D) or 1 µmol m^−2^ s^−1^ far-red light (FR) for 3 h (E-G). HA-tagged SPA proteins were detected by an α-HA antibody. HSC70 protein detected by specific antibodies served as loading control. Signals were quantified and HA/HSC70 ratios, normalized to the respective darkness controls, are indicated below the immunoblots.

We analyzed the respective SPA-HA protein level in dark-grown and light-exposed seedlings. Figure 1B confirms for the controls that exposure of dark-grown seedlings to 3 h of red light reduced SPA2-HA protein levels to almost undetectable levels, as we had reported previously (Balcerowicz *et al*., 2011; Chen *et al*., 2015), while SPA3-HA protein levels were retained in red light in three independent transgenic lines. Expression of the chimeric DS_233-HA protein resulted in its accumulation under dark conditions, whereas its abundance was markedly reduced upon exposure to red light (Figure 1C). This indicates that the N-terminal domain of SPA2 conferred instability to the C-terminal domains of SPA3. In the vice versa arrangement of the domains, the DS_322-HA protein accumulated to similar levels in red- and dark-grown seedlings (Figure 1D), indicating that the N-terminal domain of SPA3 conferred stability to the C-terminal domains of SPA2. Taken together, these results demonstrate that the N-terminal domains of SPA2 and SPA3 carry information for instability or stability, respectively, in red-exposed seedlings.

Similar results were obtained when dark-grown seedlings were exposed to FR or blue light. SPA2-HA levels were strongly reduced after short treatment with FR and B when compared to darkness, while SPA3-HA levels were similar in darkness and blue light or only slightly reduced by FR exposure (Figure S1A, Figure 1E). DS_233-HA protein levels were strongly reduced after exposure to FR or blue light in most transgenic lines (Figure 1F, Figure S1B), while DS_322-HA proteins clearly accumulated in both FR- and blue light-exposed seedlings (Figure 1G, Figure S1A). Thus, also in blue and red light, the N-terminal domain determined the degree of SPA protein stability, with that of SPA2 causing SPA degradation in response to red and blue light, and that of SPA3 conferring stability in red and blue light.

### The C-terminal domains of SPA2 and SPA3 differ in their activity in repression of photomorphogenesis

We determined the phenotype of the transgenic *spa123* lines to assess the capacity of the transgenes to complement the *spa123* mutant phenotype. *spa123* mutant seedlings exhibit constitutive photomorphogenesis in darkness, showing a short hypocotyl and open cotyledons (Figure 2A) (Laubinger *et al*., 2004). Transgenic lines expressing any of the four transgenes displayed a long hypocotyl and closed cotyledons in darkness, similar to wild-type seedlings (Figure 2A). Thus, all four transgenes complemented the *spa123* mutant phenotype in dark-grown seedlings. This finding demonstrates that SPA-HA proteins, including the chimeric DS_SPA2/SPA3 proteins, are functional.

**Figure 2.**
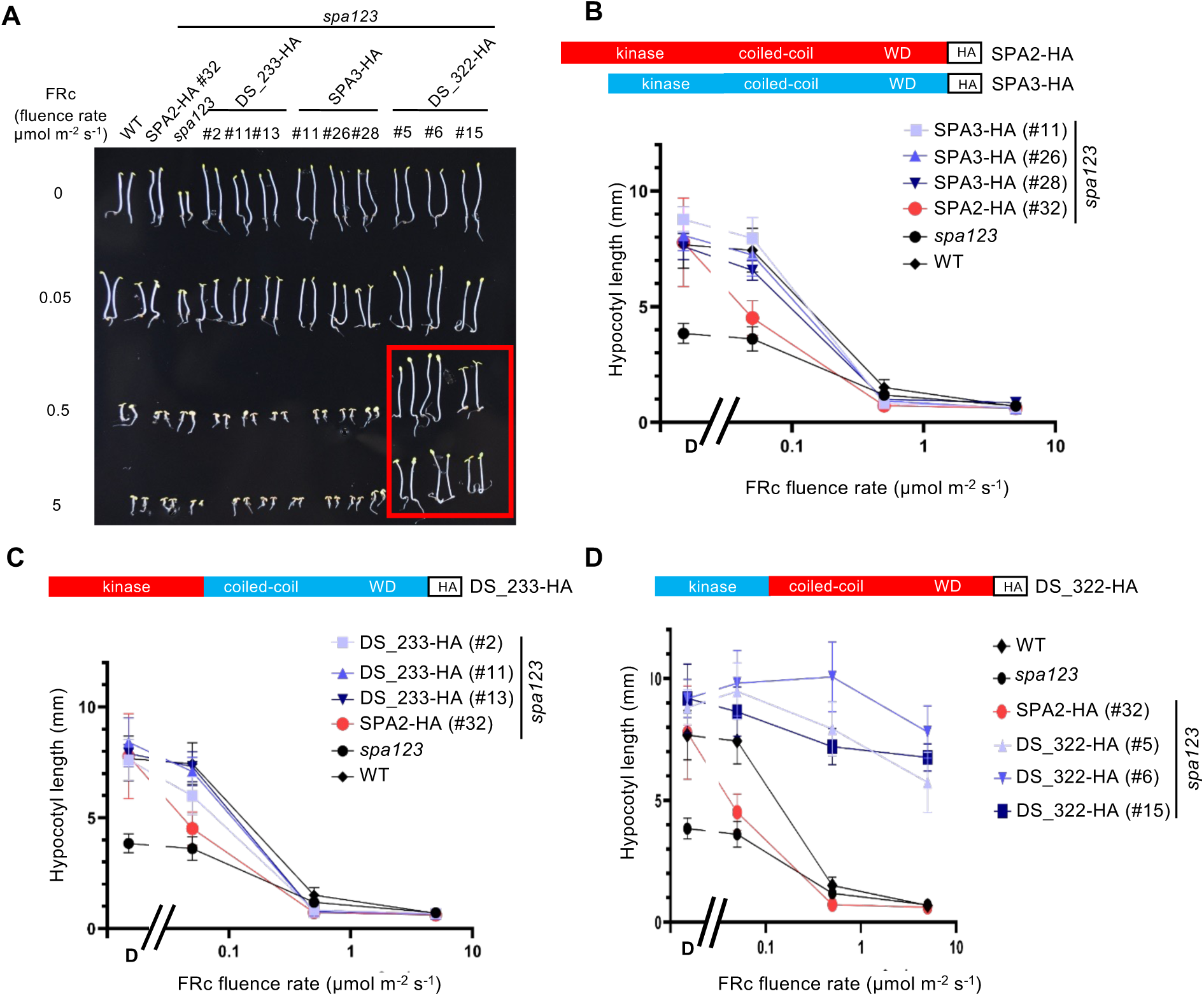
Seedlings expressing the chimeric DS_322-HA protein are insensitive to FR. A. Phenotype of 4-day-old *spa123* mutant seedlings expressing the indicated domain swap constructs described in Figure 1A. Seedlings were grown in darkness or continuous far-red light (FRc) of the indicated fluence rates for 4 days. Seedlings expressing SPA2-HA or SPA3-HA served as controls. Line numbers indicate independent transgenic lines. B-D. Quantification of hypocotyl length of *spa123* mutant seedlings expressing SPA2-HA or SPA3-HA (B), DS_233-HA (C) or DS_322-HA (D) grown as in A. Error bars indicate the SEM. Red colors in the schemes represent domains from SPA2, blue colors represent SPA3 domains.

In FR, two distinct phenotypes were observed: *spa123* lines expressing SPA2-HA, SPA3-HA or DS_233-HA exhibited a shortening of hypocotyls and an opening of cotyledons with increasing FR fluence rate in a manner similar to wild-type seedlings (Figure 2A). A quantification of the hypocotyl length confirmed the visual phenotype (Figure 2B, C). In contrast, *spa123* lines expressing DS_322-HA clearly stood out by being very insensitive to FR, showing a long hypocotyl even at the highest FR fluence rate used (Figure 2A, D). Thus, combining the N-terminal domain of SPA3 with the C-terminal domains of SPA2 generates a repressor that is barely inactivated by FR and thereby prevents photomorphogenesis in FR.

To assess whether the observed phenotypes in continuous FR (FRc) correlate with the respective SPA protein levels, we determined SPA-HA protein levels in seedlings grown in FRc for 4 days (Figure 3). As expected, SPA2-HA protein barely accumulated, while SPA3-HA protein levels were clearly detectable and even very high in two out of three transgenic lines. DS_233-HA protein levels were low in all three lines. DS_322 levels varied among the three lines.

**Figure 3.**
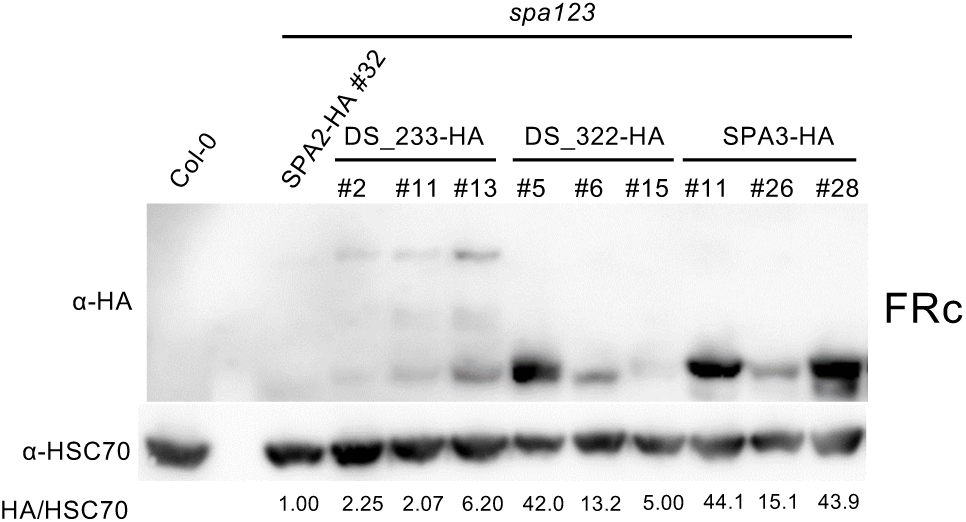
Comparative immunodetection of chimeric SPA-HA proteins in seedlings grown in FRc for 4 days. Seedlings of the indicated genotypes were grown in 5 µmol m^−2^ s^−1^ FRc for 4 days. HA-tagged SPA proteins were detected by an α-HA antibody. HSC70 protein detected by specific antibodies were used as loading control. Signals were quantified and HA/HSC70 ratios, normalized to the respective SPA2-HA/HSC70 levels, are indicated below the immunoblots.

When comparing respective SPA-HA protein levels and phenotypes in the transgenic lines (Table 1) we conclude the following: 1) it is evident that SPA2-HA lost the ability to repress photomorphogenesis in FR due to SPA2-HA protein degradation in FR. 2) The C-terminal domains of SPA2 confer much higher repressor activity than those of SPA3. This is based on the finding that DS_322-HA-expressing lines exhibited much taller hypocotyls in FR than SPA3-HA-expressing lines (Figure 2A,B,D), even when comparing lines with similar SPA-HA protein accumulation (Figure 3; DS_322-HA #5 with SPA3-HA #11 and #28; DS_322-HA #6 with SPA3-HA #26). All three DS_322 lines exhibited similar insensitivity to FR at the fluence rates used (Figure 2A,B,D) despite considerable difference in DS_322-HA protein accumulation (Figure 3). This suggests that even an only moderate stabilization of a SPA2 protein in FR leads to severe suppression of photomorphogenesis. This confirms that degradation of SPA2 in FR is essential to allow photomorphogenesis to proceed, as we found for a stable SPA2 deletion protein that lacks the N-terminal domain (Schenk *et al*., 2021). In summary, SPA2 and SPA3 do not only differ in their protein stability but also in their repressor activity – and the difference in repressor activity per se is independent of the protein stability.

**Table 1.** Activities of SPA2, SPA3 and chimeric DS_SPA domain-swap proteins in repression of photomorphogenesis in FRc.

| SPA-HA protein expressed in transgenic seedlings | SPA-HA protein level in FRc | Hypocotyl length in FRc | SPA-HA repressor activity in FRc |
| --- | --- | --- | --- |
| SPA2-HA | very low | short | low |
| SPA3-HA | high | short | low |
| DS_233-HA | low | short | low |
| DS_322-HA | intermediate-high | tall | very high |

We also analyzed the phenotype of these transgenic lines when seedlings were grown in red or blue light. The phenotypes in blue light resembled those described above for FRc: seedlings expressing the light-stable DS_322-HA fusion protein were insensitive to blue light, while lines expressing SPA2-HA, SPA3-HA or DS_233-HA were responsive to blue light similar to wild-type seedlings (Figure S2A-D). Because DS_322-HA and SPA3-HA levels were similar in blue light (Figure S2E), these results indicate again that the C-terminal domains of SPA2 in DS_322-HA confer much higher activity in repression of photomorphogenesis in blue light than those of SPA3 in SPA3-HA.

In red light, similar observations were made, though less pronounced (Figure S3A-E). DS_322 expressing seedlings were less responsive to red light than those expressing SPA3-HA, despite accumulating similar SPA-HA protein levels. However, 322_HA lines were not as insensitive to red light as they were to FR and blue light (Figure S3D, S2D, 2D).

### Differential interaction of the N-terminal domains of SPA2 and SPA3 with phyA

We previously defined that phytochromes, in particular phyA, mediate SPA2 degradation in red, FR and blue light, while cryptochromes do not have a major role in this process, not even in blue light (Chen *et al*., 2015). We therefore asked whether SPA2 and SPA3 differ in their interaction with phyA. Since the N-terminal domains of SPA2 and SPA3 confer SPA protein stability information we investigated the interaction of these domains with phyA. In a yeast two-hybrid assay, the N-terminal domain of SPA2 (SPA2-NT) strongly interacted with phyA in a red light-dependent fashion (Figure 4A). In contrast, SPA3-NT did not interact with phyA (Figure 4A). In this assay, we included also SPA4 which is closely related in sequence to SPA3 and also a light-stable protein (Schenk *et al*., 2021). SPA4-NT also did not interact with phyA.

**Figure 4.**
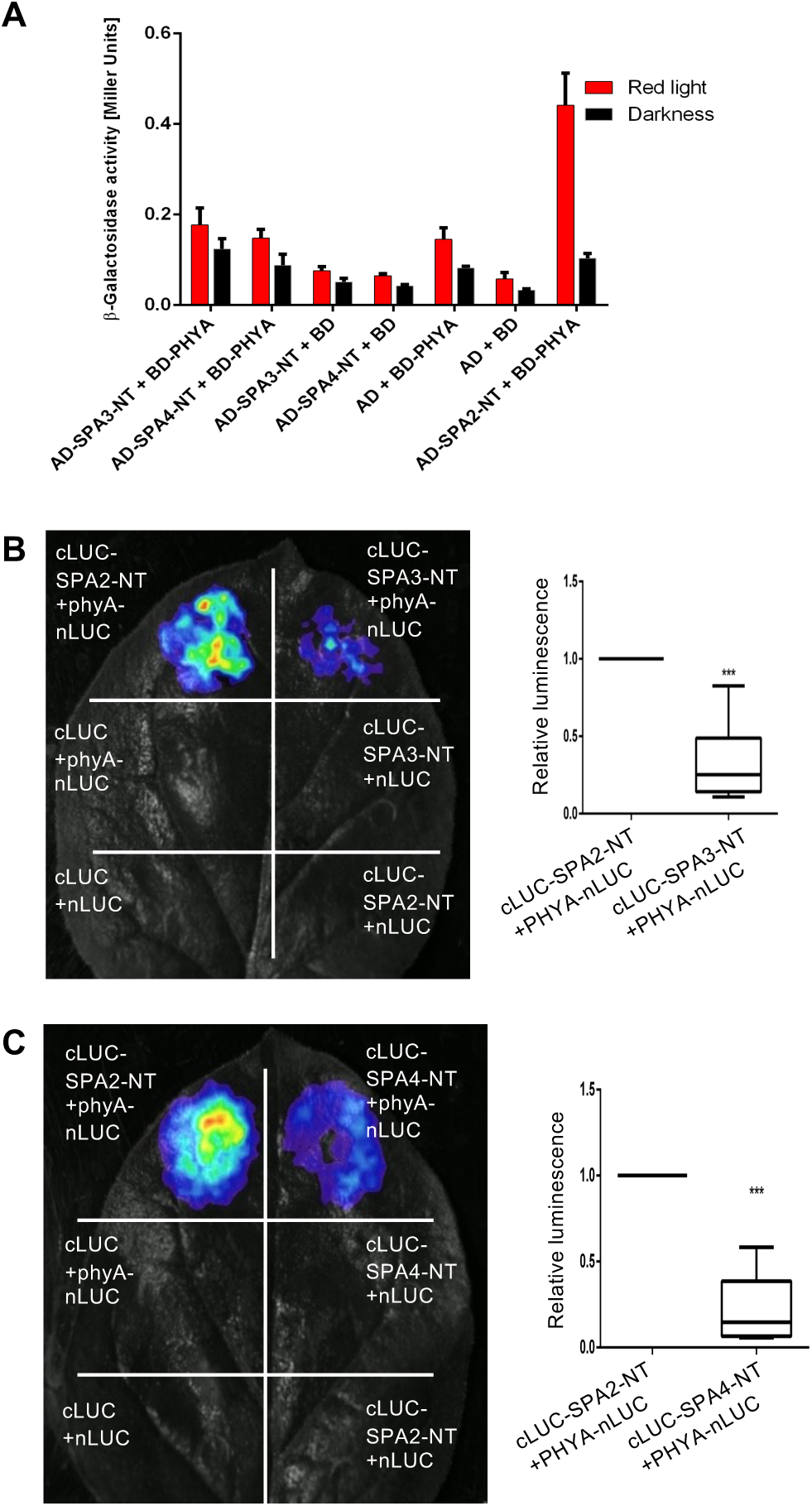
The N-terminal domains of SPA3 and SPA4 barely interact with phyA. A. Yeast two hybrid assay. The indicated proteins fused to the GAL4 transcription activation domain (AD) or DNA-binding domain (BD) were co-expressed in yeast exposed to darkness or red light (1 µmol m^−2^ s^−1^) for 2 days. B,C. Luciferase complementation imaging assays in transfected *N. benthamiana* leaves investigating the interaction between SPA3-NT (B) or SPA4-NT (C) with phyA. SPA2/3/4-NT were fused to the C-terminal half of luciferase (cLUC), phyA was fused to the N-terminal half of luciferase (nLUC). All proteins contained an artificial nuclear localization sequence to ensure nuclear import. (left) Representative images of luciferase activity in *N. benthamiana* leaves three days after infiltration with the indicated combinations of constructs. (Right) Quantification of relative luciferase activity (n=6 leaves). The upper and lower hinges of the box plot correspond to the first and third quartiles, respectively. Whiskers extend from the hinges to the highest/lowest value within the 1.5 * interquartile range (IQR). The median is represented by a horizontal line within the box. Asterisks indicate a significant difference in the relative luminescence signal analyzed using student’s t-test with *** p < 0.001.

To corroborate these findings in planta we performed luciferase complementation imaging (LCI) after expressing split luciferase constructs in transfected tobacco leaves. To ensure nuclear import, all proteins were fused to an artificial nuclear localization sequence (NLS). In these assays, SPA2-NT interacted much more strongly with phyA than SPA3-NT or SPA4-NT (Figure 4B, C). Thus, the ability of phyA to mediate SPA2 degradation in the light correlates with its ability to interact with the N-terminal domain of SPA2.

### phyA interacts with several regions in the N-terminal domain of SPA2

A sequence comparison between the N-terminal domains of SPA2 and SPA3 reveals that SPA2 carries an N-terminal extension not found in SPA3 (Figure 5A, named SPA2-NT-1). The middle region of SPA2 (SPA2-NT-2) carries very low sequence identity with SPA3, while the third region carries considerable sequence identity with SPA2 (Figure 5A) (Laubinger and Hoecker, 2003). We hypothesized that the regions with lowest sequence conservation between SPA2 and SPA3 may be responsible for the specific interaction of SPA2-NT with phyA. We therefore investigated the interaction of phyA with each of the three regions of SPA2-NT. In yeast two-hybrid we found that SPA2-NT-1 displayed a similarly strong, red-light dependent interaction with phyA as SPA2-NT did (Figure 5B). SPA2-NT-2 interacted weakly with phyA, while SPA2-NT-3 did not interact with phyA (Figure 5B). In LCI assays, all three regions, fused to an artificial NLS, interacted with phyA and these interactions were at least as strong as that with SPA2-NT (Figure 5C-F). This suggests that phyA interacts with multiple regions within the N-terminal domain of SPA2.

**Figure 5.**
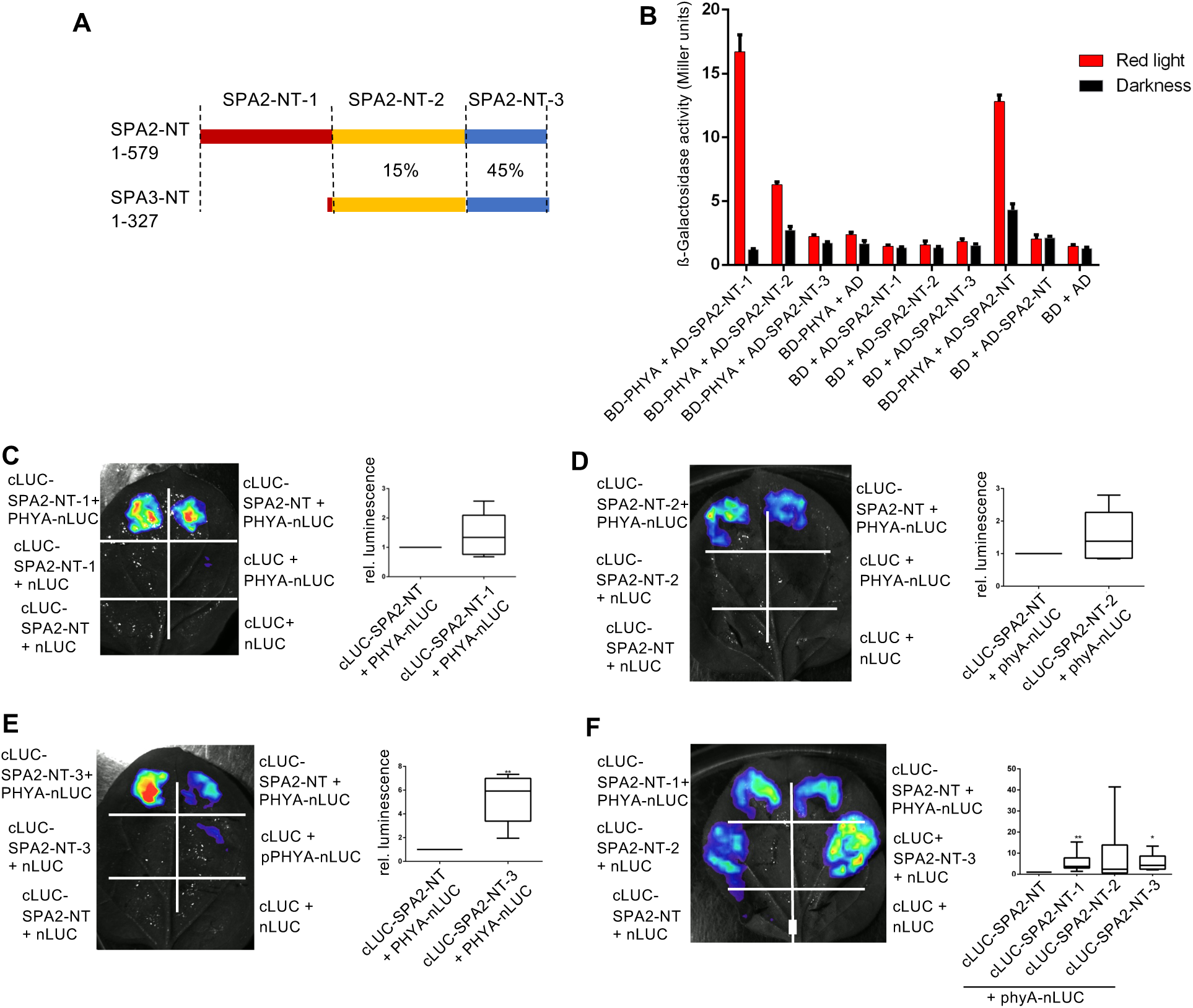
phyA interacts with multiple regions of SPA2-NT. A. Schematic representation of SPA2-NT and SPA3-NT regions and the amino acid sequence identity. B. Yeast two-hybrid assay showing that SPA2-NT-1, SPA2-NT-2 and SPA2-NT interact with phyA in red light. The indicated proteins fused to the GAL4 transcription activation domain (AD) or DNA-binding domain (BD) were co-expressed in yeast exposed to darkness or red light (1 µmol m^−2^ s^−1^) for 2 days. NT-1: amino acids 1-234 of SPA2, NT-2: 235-461 of SPA2; NT-3: 462-579 of SPA2. Error bars indicate the SEM (n=3). C-F. Luciferase complementation imaging assays in transfected *N. benthamiana* leaves investigating the interaction between phyA and SPA2-NT-1 (C), SPA2-NT-2 (D), SPA2-NT-3 (E) and all three SPA2-NTs (F). All SPA2 deletion proteins were fused to an artificial nuclear localization sequence to ensure nuclear import. Statistical analysis as described in Fig. 4B,C (n=6 leaves).

## Discussion

In a complex with COP1, SPA proteins are key repressors of photomorphogenesis that allow plants to etiolate in darkness, to exhibit a shade avoidance response and to suppress flowering under non-inductive short day photoperiods (Ponnu and Hoecker, 2021; Zhou and Deng, 2025). While COP1 is a single copy gene in most angiosperms, there are multiple *SPA* genes, e.g. four *SPA* genes in Arabidopsis, two *SPA* genes in rice and 10 *SPA* genes in soybean (Laubinger *et al*., 2004; Li *et al*., 2023; Qin *et al*., 2023; Ranjan *et al*., 2014). Phenotypic analysis of Arabidopsis *spa* mutants has shown that the four *SPA* genes have overlapping but also partially distinct functions during growth and development (Menon *et al*., 2016). Here, we addressed the functional divergence of SPA2 and SPA3 which represent different subclades of the four-member *SPA* gene family (Laubinger and Hoecker, 2003; Ranjan *et al*., 2014).

Light triggers the degradation of SPA1 and, notably, SPA2, which becomes highly unstable when seedlings are exposed to even a few minutes of very low-fluence light (Balcerowicz *et al*., 2011; Chen *et al*., 2015). The light-dependent degradation of SPA2 is essential for photomorphogenesis, as SPA2 mutations that stabilize the SPA2 protein render seedlings insensitive to light (Schenk *et al*., 2021). In contrast, SPA3 and SPA4 proteins remain stable under light conditions (Schenk *et al*., 2021). Here, we have shown that the distinct protein stabilities of SPA2 and SPA3 are determined by their respective N-terminal domains. Furthermore, we have demonstrated that the C-terminal domains of SPA2 and SPA3 differ in their abilities to repress photomorphogenesis in light-grown seedlings. Together, these findings indicate that SPA2 and SPA3 have undergone substantial functional divergence since their gene duplication event.

The domain-swap analyses between SPA2 and SPA3 domains indicate that the respective N-terminal domains confer the information for the distinct protein stability of SPA2 and SPA3 in light-exposed seedlings: The N-terminal domain of SPA2 confers protein degradation in response to light, while the N-terminal domain of SPA3 maintains SPA protein stability after exposure to light (Figure 6). This is consistent with previous findings showing that deletion-derivatives of SPA1 and SPA2 that lack the respective N-terminal domain are not degraded in response to light (Fittinghoff *et al*., 2006; Schenk *et al*., 2021; Yang and Wang, 2006). Moreover, the greatly reduced stability of SPA2 when compared to SPA1 also mapped to the respective N-terminal domain (Chen *et al*., 2016).

**Figure 6.**
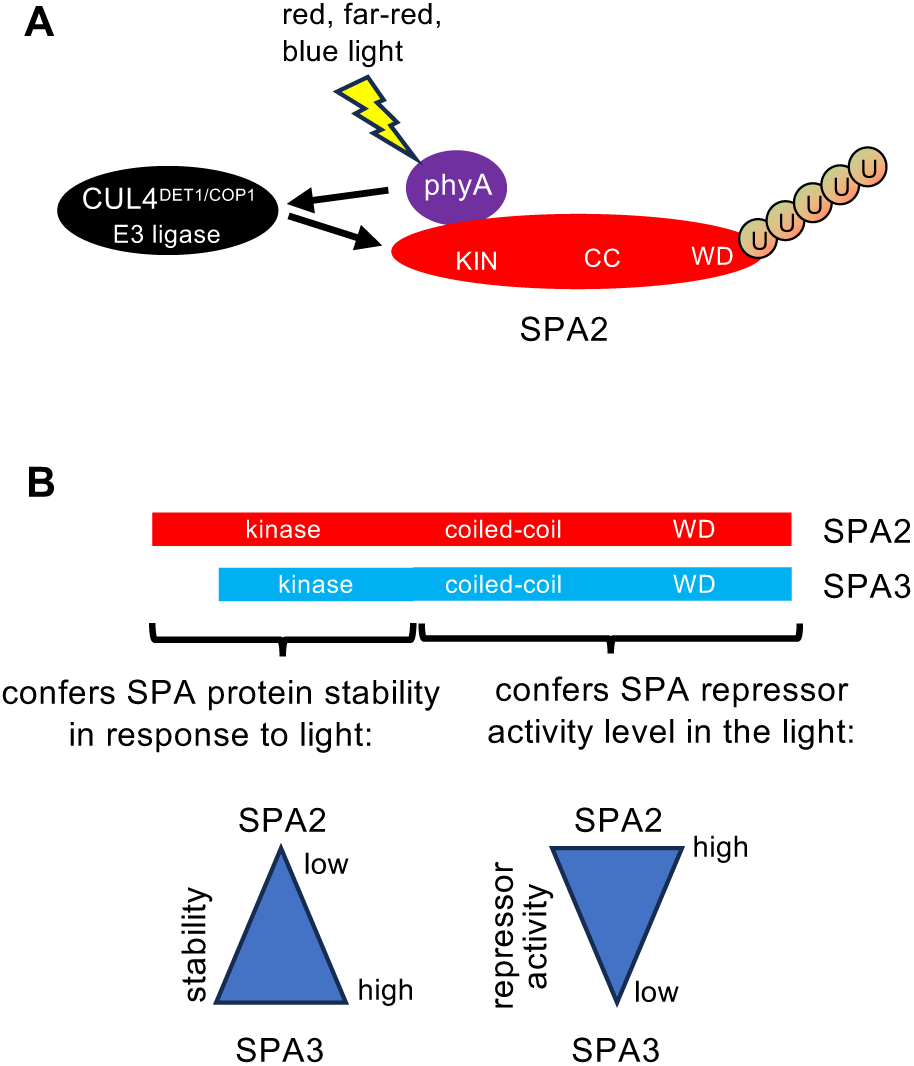
Model on the divergent activities of SPA2 and SPA3 domains in light-grown seedlings. A. Interaction of photo-activated phyA with the N-terminal kinase domain (KIN) of SPA2 confers SPA2 degradation via the CUL4^DET1/COP1^ E3 ubiquitin ligase. B. Differential functions of the N-terminal kinase domain and C-terminal coiled-coil and WD-repeat domains of SPA2 and SPA3: The SPA N-terminal, kinase-containing domain is responsible for the distinct stability of SPA2 and SPA3 proteins in response to light: that of SPA2 confers degradation, while the that of SPA3 confers protein stability in light-exposed seedlings. The C-terminal domains (coiled-coil and WD, CC-WD) of SPA2 and SPA3 confer a differential repressor activity level in light-grown seedlings: SPA2-CC-WD retains high repressor activity, while SPA3-CC-WD retains low repressor activity.

We have previously shown that phytochromes are responsible for the rapid degradation of SPA2 in response to light, while cryptochromes do not have a major role in this process. In particular, phyA is required for SPA2 degradation in far-red and blue light (Chen *et al*., 2015). Consistent with this finding, the N-terminal domain of SPA2 interacts with phyA, while that of SPA3 does not. This suggests that the light-induced interaction of phyA with the N-terminal domain of SPA2 causes the polyubiquitination and subsequent degradation of SPA2, while the failure of phyA to interact with the N-terminal domain of SPA3 renders SPA3 stable in light-grown seedlings (Figure 6A, B). Mechanistically, phyA-binding to the N-terminal domain of SPA2 might promote the association of SPA2 with the CUL4^DET1/COP1^-complex which is responsible for SPA2 polyubiquitination (Figure 6A) (Chen *et al*., 2015; Schenk *et al*., 2021). Interestingly, the CUL4-DET1 complex was also shown to destabilize COP1 in plants and humans, though in a light-independent manner in plants (Burgess *et al*., 2025; Cañibano *et al*., 2021). phyA-binding to the N-terminal, kinase domain-containing region of SPA2 might also alter the kinase activity of SPA2 which may subsequently allow CUL4^DET1/COP1^ polyubiquitination of SPA2. However, missense mutations in the SPA1 kinase domain that strongly reduce the kinase activity of SPA1 (Paik *et al*., 2019) did not alter SPA1 protein levels in light-grown plants (Holtkotte *et al*., 2016). Beyond that, we do not exclude the possibility that lysine residues that are polyubiquitinated in SPA2 are not conserved in SPA3, thereby rendering SPA3 stable.

Our domain swap analysis also uncovered that the C-terminal domains of SPA2 and SPA3 greatly differ in their repressor activity strength, with those of SPA2 retaining much stronger repressor activity in the light when compared to those of SPA3 (Figure 6B). This becomes evident in the context of a light-stable hybrid SPA protein, i.e. when the C-terminal domains of SPA2 are fused to the protein-stabilizing N-terminal domains of SPA3: these 322-expressing transgenic seedlings were insensitive to light, exhibiting a long hypocotyl even at high fluence rates of light. Mechanistically, the quantitative difference between the repressor activities of the respective C-terminal domains (CC+WD) in the light may reflect the effectiveness of light-induced COP1/SPA inactivation by photoreceptors. For example, since photoreceptors disrupt the COP1/SPA interaction which is formed through the CC domains, the CC domain of SPA2 may retain a stronger interaction with COP1 in light-grown seedlings (Hoecker and Quail, 2001; Lian *et al*., 2011; Liu *et al*., 2011; Lu *et al*., 2015; Sheerin *et al*., 2015). Alternatively, photoreceptors may be less effective to competitively displace substrates from binding to the WD domains in a COP1/SPA2 complex than in a COP1/SPA3 complex (Lau *et al*., 2019; Ponnu *et al*., 2019; Trimborn *et al*., 2025). Since the light-induced shuttling of COP1 from the nucleus to the cytosol is SPA-dependent (Balcerowicz *et al*., 2017), it is also possible that the C-terminal domains of SPA2 retain a stronger nuclear localization of COP1 than those of SPA3. Though we detect the differential repressor activities of SPA2 and SPA3 CC+WD domains only in light-grown seedlings using the domain-swap set up, we do not exclude the possibility that COP1/SPA2 complexes are generally more effective repressors of photomorphogenesis than COP1/SPA3 complexes, i.e. also in darkness. Consistent with this idea, dark-grown *spa134* triple mutants (retaining only SPA2 function) fully etiolate in darkness, while *spa124* triple mutants (retaining only SPA3 function) undergo constitutive photomorphogenesis, indicating lower repressor activity of SPA3 than of SPA2 (Laubinger *et al*., 2004).

Taken together, our domain swap approach demonstrates that the domains of SPA2 and SPA3 have undergone considerable functional divergence during evolution. It is interesting that the SPA2 protein combines an N-terminal instability domain with a strong repressor domain (CC+WD). The evolution of a light-induced instability of SPA2 was likely necessary to counteract the strong repressor activity of SPA2 and to, thereby, allow rapid inactivation of COP1/SPA2 as germinating seedlings emerge from soil to the light. This mechanism facilitates rapid de-etiolation and the initiation of photosynthesis which is essential for seedling survival. Thus, these properties of the SPA2 protein enable it to function as a most potent repressor of photomorphogenesis in darkness, as evident from the full etiolation of dark-grown *spa134* seedlings, while exposure to light allows very rapid deetiolation, as observed in light-grown *spa134* seedlings that resemble *spaQ* quadruple mutant seedlings (Laubinger *et al*., 2004). SPA3, in contrast, has a most pronounced role in adult plants rather than in seedlings (Laubinger and Hoecker, 2003). Here, the SPA3-mediated promotion of leaf and petiole expansion may be sufficiently light-regulated by a SPA protein having the features of SPA3 which are likely shared by the very closely related SPA4 protein.

## Supplementary data

**Figure S1.**
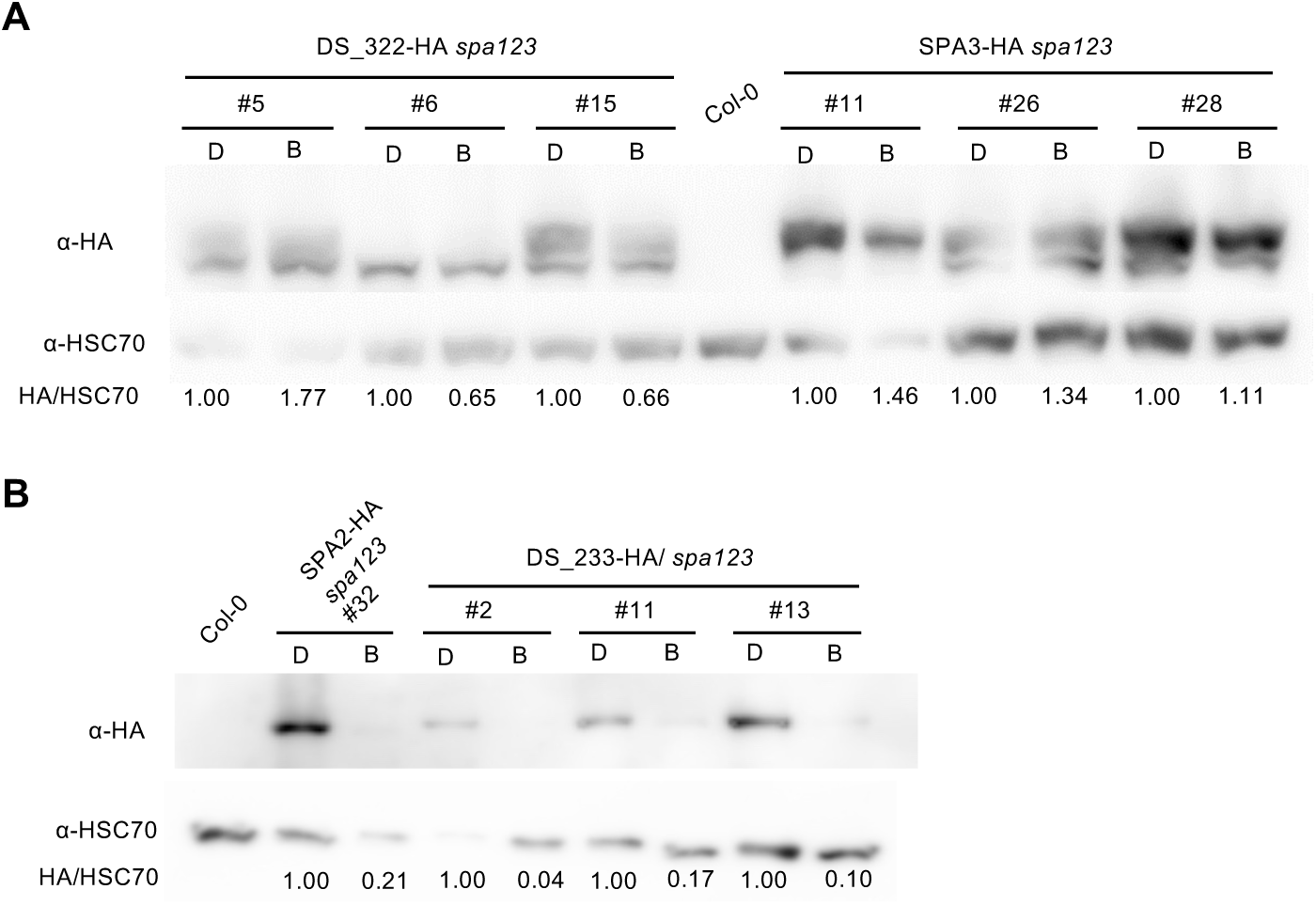
The N-terminal domains of SPA2 and SPA3 confer SPA protein stabilty information in blue light. A, B. Immunodetection of the indicated HA-tagged SPA proteins in *spa123 triple* mutant seedlings grown in darkness for 4 days (D) followed by transfer to 5 µmol m^−2^ s^−1^ blue light (B) for 3 h. HA-tagged SPA proteins were detected by an α-HA antibody. HSC70 protein detected by specific antibodies served as loading control. Signals were quantified and HA/HSC70 levels, normalized to the respective darkness controls, are indicated below the immunoblots.

**Figure S2.**
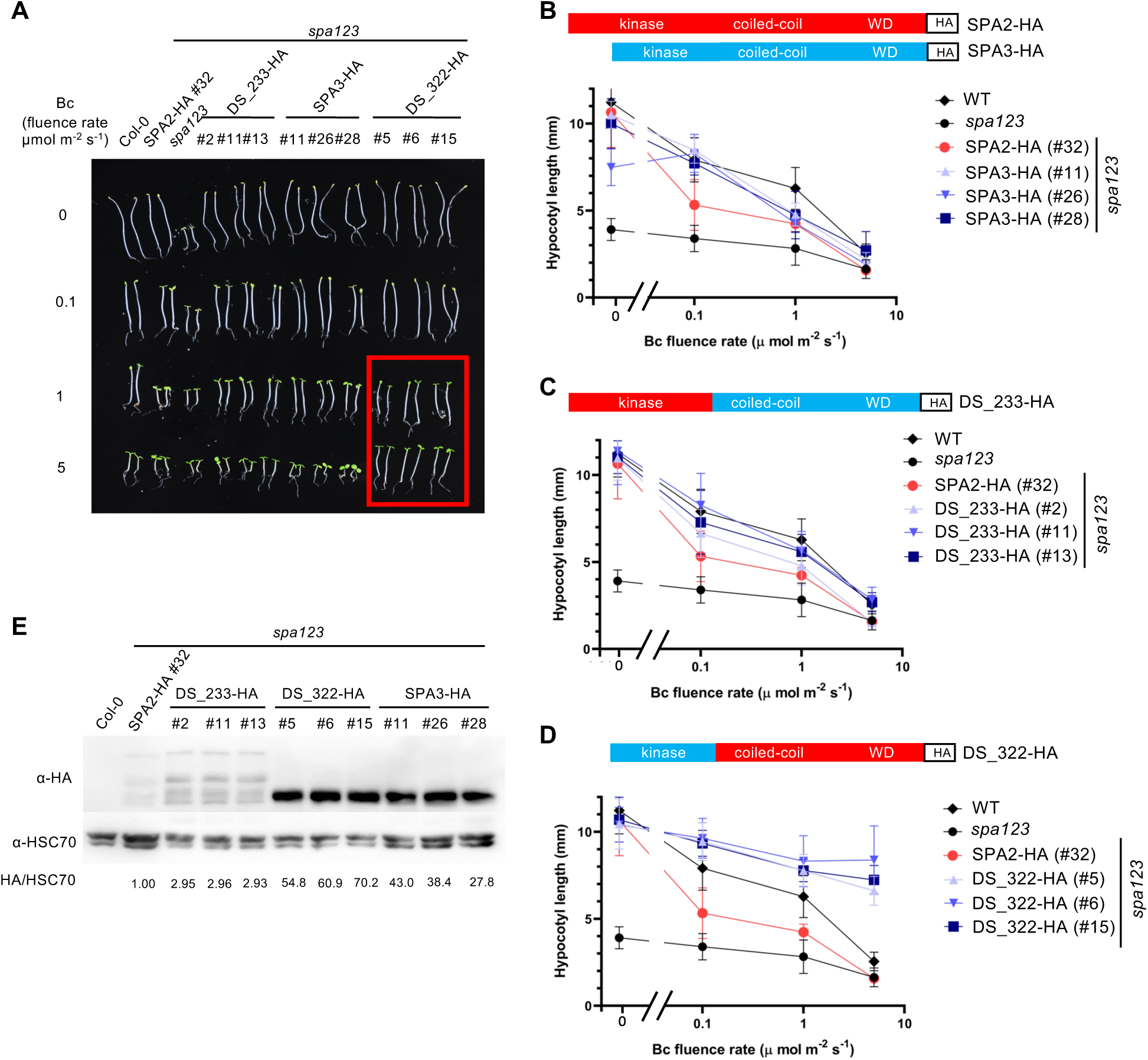
Seedlings expressing the chimeric DS_322-HA protein are insensitive to blue light. A. Phenotype of 4-day-old *spa123* mutant seedlings expressing the indicated domain swap constructs described in Figure 1. Seedlings were grown in darkness or continuous blue light (Bc) of the indicated fluence rates. Seedlings expressing SPA2-HA or SPA3-HA served as controls. Line numbers indicate independent transgenic lines. B-D. Quantification of hypocotyl length of *spa123* mutant seedlings expressing SPA2-HA or SPA3-HA (B), DS_233-HA (C) or DS_322-HA (D) grown as in A. Error bars indicate the SEM. Red colors in the schemes represent domains from SPA2, blue colors represent SPA3 domains. E. SPA-HA protein levels in the indicated genotypes grown in 10 µmol m^−2^ s^−1^ Bc for 4 days. HA-tagged SPA proteins were detected by an α-HA antibody. HSC70 protein detected by specific antibodies were used as loading control. Signals were quantified and HA/HSC70 ratios, normalized to the respective SPA2-HA/HSC70 levels, are ndicated below the immunoblots.

**Figure S3.**
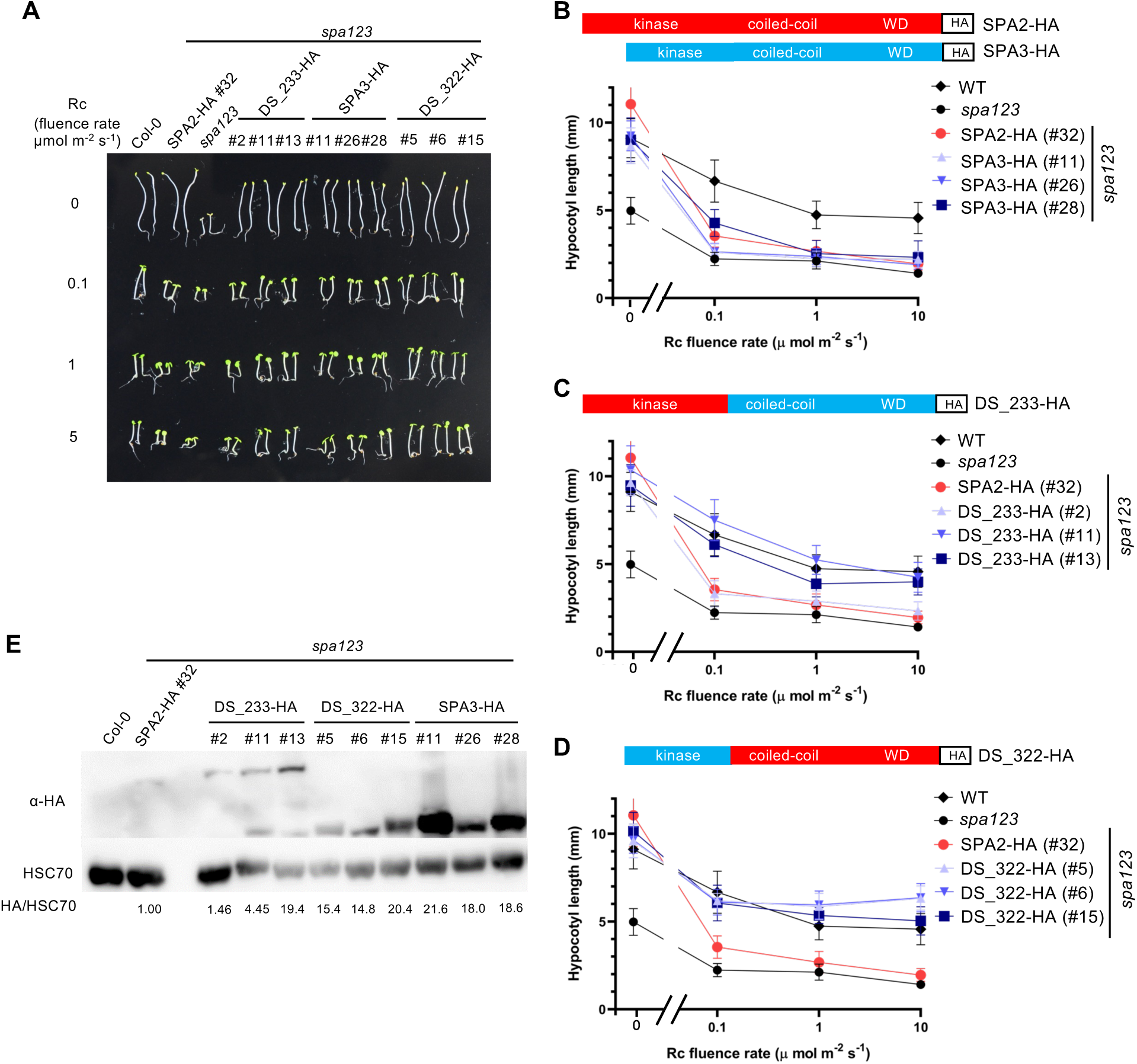
Seedlings expressing the chimeric DS_322-HA protein exhibit a reduced sensitivity to red light. A. Phenotype of 4-day-old *spa123* mutant seedlings expressing the indicated domain swap constructs described in Figure 1. Seedlings were grown in darkness or continuous red light (Rc) of the indicated fluence rates. Seedlings expressing SPA2-HA or SPA3-HA served as controls. Line numbers indicate independent transgenic lines. B-D. Quantification of hypocotyl length of *spa123* mutant seedlings expressing SPA2-HA or SPA3-HA (B), DS_233-HA (C) or DS_322-HA (D) grown as in A. Error bars indicate the SEM. Red colors in the schemes represent domains from SPA2, blue colors represent SPA3 domains. E. SPA-HA protein levels in the indicated genotypes grown in 10 µmol m^−2^ s^−1^ Rc for 4 days. HA-tagged SPA proteins were detected by an α-HA antibody. HSC70 protein detected by specific antibodies were used as loading control. Signals were quantified and HA/HSC70 ratios, normalized to the respective SPA2-HA/HSC70 levels, are indicated below the immunoblots.

## Acknowledgements

We thank the greenhouse staff and many undergraduate students for expert care of our plants. We are grateful to Andreas Hiltbrunner for the BD-phyA plasmid.

## Author contributions

UH and ZC designed the experiments. ZC and VF performed the experiments. All authors analyzed the data. ZC and UH discussed the results and wrote the manuscript.

## Conflict of interest

The authors declare no conflict of interest.

## Funding

This work was supported by a grant from the Deutsche Forschungsgemeinschaft (HO 2793/3-5 to U.H.) and Germanýs Excellence Strategy EXC 2048/1 Project ID: 390686111 (to U.H.). Z.C. was a recipient of a stipend from the Chinese Scholarship Council.

## Data availability

All data supporting the findings of the study are available within the paper and within its supplementary data published online.

## Supplemental Materials and Methods

### Generation of plasmid constructs

For all primer sequences, see Table S1.

#### SPA2-SPA3 domain swaps

For the domain swap, a sequence conserved between SPA2 and SPA3 was used; it encodes the amino acid sequence LQSE (SPA2 amino acids 545-548; SPA3 amino acid 292-295) and is located between the respective kinase domain and the coiled-coil domain. First, the pJHA-SPA3-HA plasmid expressing SPA3-3xHA under the control of the native *SPA2* 5’ and 3’ regulatory sequences (Schenk *et al*., 2021) was used as a backbone and digested with ApaI and MluI restriction enzymes to remove the *SPA3* open-reading-frame and part of the 3xHA tag (the MluI site is located within the 3xHA tag).

To generate the domain swap DS-233-HA construct expressing the N-terminal domain of SPA2 and the coiled-coil and WD-repeat domains of SPA3 fused to a 3xHA tag under the control of the *SPA2* 5’and 3’regulatory sequences, the N-terminal sequence of SPA2 was PCR-amplified from a SPA2-3xHA-expressing plasmid (Balcerowicz *et al*., 2017; Schenk *et al*., 2021) as a template using the primers ol-2656 and ol-2657. The sequence encoding the C-terminal coiled-coil and WD-repeat domains of SPA3 were amplified from a SPA3-3xHA encoding plasmid using the primers ol-2658 and ol-2659. The primer ol-2659 contained the complementary sequence of the 3xHA tag, which matched the backbone that had been digested ApaI and MluI. All four primers contained an overhang for subsequent overhang extension PCR. Both PCR products were purified from agarose gels and used as templates to generate the 233 domain fusion by overhang extension PCR using the primers ol-2656 and ol-2659. After gel purification, the 233 fusion sequence was cloned into the ApaI/MluI-digested backbone plasmid via ClonExpress II One Step Cloning Kit (Absource Diagnostics GmbH, Munich, Germany).

To generate the domain swap DS-322-HA construct expressing the N-terminal domain of SPA3 and the coiled-coil and WD-repeat domains of SPA2 fused to a 3xHA tag, the N-terminal sequence of SPA3 was PCR-amplified from a SPA3-3xHA-expressing plasmid (Schenk *et al*., 2021) as a template using the primers ol-2660 and ol-2661. The sequence encoding the C-terminal coiled-coil and WD-repeat domains of SPA2 were amplified from a SPA2-3xHA encoding plasmid using the primers ol-2662 and ol-2663. Both PCR products were purified from agarose gels and used as templates to generate the 233 domain fusion by overhang extension PCR using the primers ol-2660 and ol-2663. After gel purification, the 233 fusion sequence was cloned into the ApaI/MluI-digested backbone plasmid as described above.

The SPA2-HA construct (Balcerowicz *et al*., 2011) and SPA3-HA construct (Schenk *et al*., 2021) were described previously, there named SPA2::SPA2-HA and SPA2::SPA3-HA, respectively.

#### Gateway Entry clones

Entry clones were generated by digesting pENTR3C with BamHI and XhoI followed by insertion of respective PCR-amplified fragments carrying overlapping ends using the ClonExpress II One Step Cloning Kit (Absource Diagnostics GmbH, Munich, Germany). To generate SPA2-NT-1, NT-2 and NT-3 entry clones the primer pairs ol-3001 and ol-3002, ol-3003 and ol-3004, and ol-3005 and ol-3006 were used, respectively. To generate pENTR3C_SPA3-NT (coding for amino acids 1-327 of SPA3) the primers ol-2125 and ol-2126 were used. To generate pENTR3C_SPA4-NT (coding for amino acids 1-298 of SPA4) the primers ol-2123 and ol-2124 were used. To generate pENTRY3C_NLS_SPA3-NT (coding for amino acids 1-327 of SPA3 fused to an N-terminal NLS) the primers ol-2806 and ol-2126 were used. To generate pENTRY3C-NLS-SPA4-NT (coding for amino acids 1-298 of SPA4 fused to an N-terminal NLS) the primers ol-2807 and ol-2124 were used.

#### Yeast two-hybrid constructs

AD-SPA2-NT was generated by Gateway cloning using the vector pACT2 that had been modified to contain a Gateway cassette (described in Schenk et al., 2021). To generate AD-SPA3-NT, AD-SPA4-NT, AD-SPA2-NT-1, AD-SPA2-NT-2 and AD-SPA2-NT-3 the above-described ENTRY clones were used to clone the respective fragments into the Gateway compatible pACT2 vector. BD-phyA (D153ah-phyA) was described in (Hiltbrunner *et al*., 2006).

#### Split-luciferase constructs

Expression vectors for split-LUC assay were generated using the ClonExpress II One Step Cloning Kit (Absource Diagnostics GmbH, Munich, Germany) or In-Fusion Snap Assembly Master Mix (Takara Bio Europe, Saint Germain-enLaye, France). The pCambia1300 nLUC and cLUC plasmids (Chen *et al*., 2008) were digested with KpnI and SalI to insert PCR-amplified fragments containing overlapping ends. The respective pENTR3C vectors (see above) were used as PCR-templates for SPA. The D153AH-PHYA plasmid (Hiltbrunner *et al*., 2006) was used to amplify the phyA fragment. To generate pCAMBIA1300-nLUC-NLS-phyA (expressing phyA-nLUC fused to an N-terminal NLS) the primers ol-2481 and ol-2482 were used. To generate pCAMBIA1300-cLUC-NLS-SPA2-NT579 expressing cLUC-SPA2-NT (amino acids 1-579 of SPA2 fused to an N-terminal NLS) primers ol-2554 and ol-2559 were used. To generate pCAMBIA1300-cLUC-NLS-SPA3-NT327 expressing cLUC-SPA3-NT (amino acids 1-327 of SPA3 fused to an N-terminal NLS) primers ol-2471 and ol-2472 were used. To generate pCAMBIA1300-cLUC-NLS-SPA4-NT298 expressing cLUC-SPA4-NT (amino acids 1-298 of SPA4 fused to an N-terminal NLS) primers ol-2473 and ol-2474 were used. To generate pCAMBIA1300-cLUC-NLS-SPA2-NT-1 expressing cLUC-SPA2-NT-1 (amino acids 1-234 of SPA2 fused to an N-terminal NLS) primers ol-2554 and ol-2997 were used. To generate pCAMBIA1300-cLUC-NLS-SPA2-NT-2 expressing cLUC-SPA2-NT-1 (amino acids 1-234 of SPA2 fused to an N-terminal NLS) primers ol-2998 and ol-2999 were used. To generate pCAMBIA1300-cLUC-NLS-SPA2-NT-3 expressing cLUC-SPA2-NT-3 (amino acids 462-579 of SPA2 fused to an N-terminal NLS) primers ol-3000 and ol-2559 were used.

---

**Table S1.**
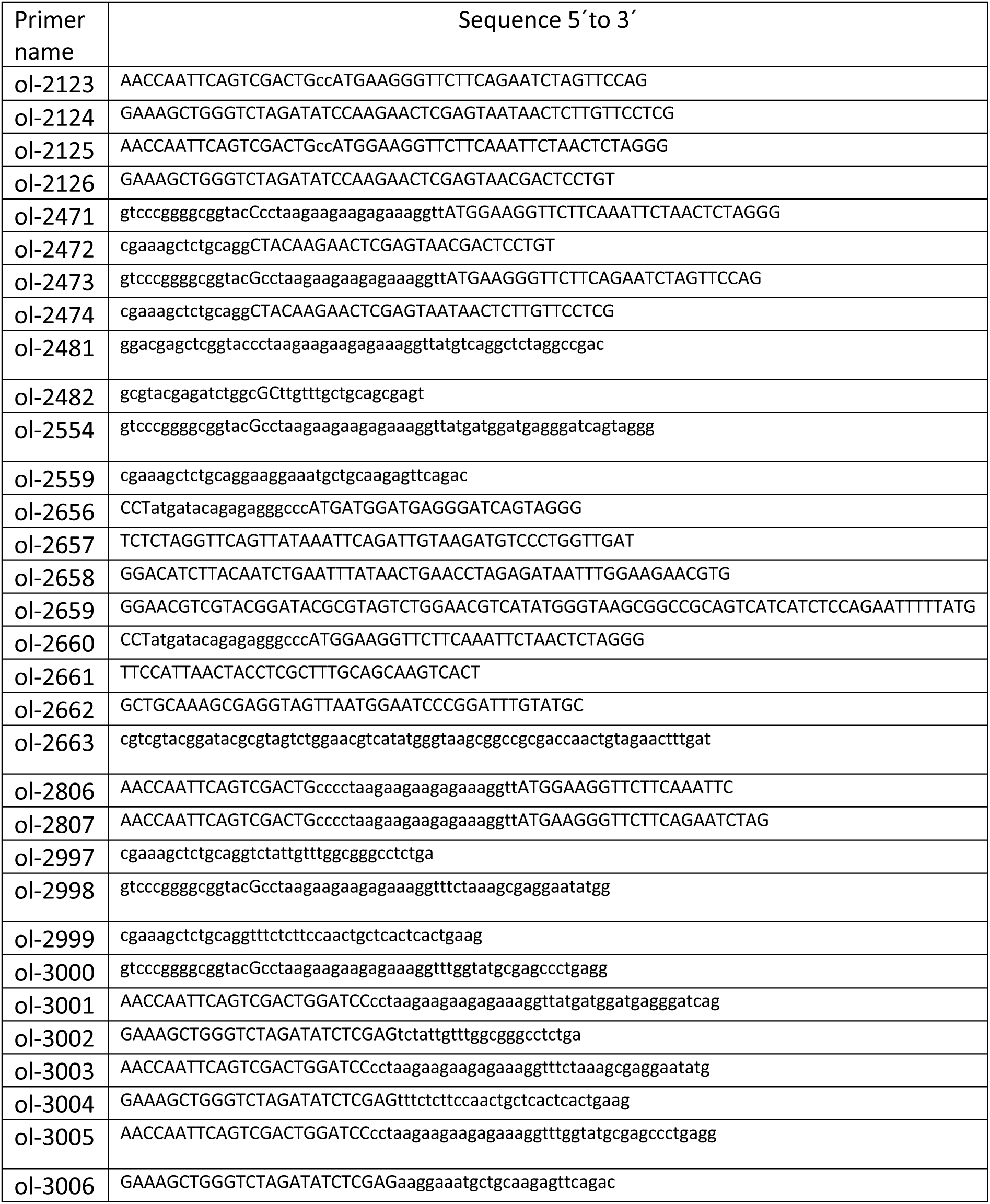
Primer sequences.

## Notes

### Competing Interest Statement

The authors have declared no competing interest.

